# Comparison analysis on transcriptomic of different human trophoblast development model

**DOI:** 10.1101/2021.02.06.430084

**Authors:** Yajun Liu, Yilin Guo, Ya Gao, Guiming Hu, Jingli Ren, Jun Ma, Jinquan Cui

## Abstract

**Aims:** Multiple models of trophoblastic cell development were developed. However, systematic comparisons of these cell models are lacking.

**Methods and Results:** In this study, first-trimester chorionic villus and decidua tissues were collected. Transcriptome data was acquired by RNA-seq and the expression levels of trophoblast specific transcription factors were identified by immunofluorescence and RNA-seq data analysis.

Differentially expressed genes between chorionic villus and decidua tissues and its related biological functions were identified. We identified genes that were relatively highly expressed and enriched transcription factors in trophoblast cells of different trophoblast cell models.

**Conclusions:** This analysis is of certain significance for further exploration of the development of placenta and the occurrence of pregnancy-related diseases in the future. The datasets and analysis provide a useful source for the researchers in the field of the maternal-fetal interface and the establishment of pregnancy.

## Introduction

Implantation failure and insufficient placental development are important causes of female infertility, recurrent miscarriage, and other pregnancy-related problems [1]. Based on different trophoblastic cell models, many molecular mechanisms for the establishment and maintenance of pregnancy have been obtained. In particular, the early stages of pregnancy have a significant impact on pregnancy outcomes [2] [3] [4]. Models of trophoblast-like cells differentiation from stem cells provide insights into the field. In previous studies, this model was compared with the transcriptome of primary cytotrophoblast recovered from term placentae trophoblastic cells ^4^. Due to Placental tissues at different stages of development are quite different, transcriptome data from villi in early pregnancy could provide further insights into this area.

In this study, we collected human first-trimester chorionic villus and decidual tissue from the same patient, performed high-throughput RNA sequencing. In particular, we analysis the expression of important trophoblastic cell-specific factors. Next, highly expressed genes in different trophoblastic cell models, including hESC line cells after BMP4 treatment (TB) comparison with H1 [6] and trophectoderm (TE) in comparison with pluripotent epiblast (EPI) cells [7], chorionic villus (CV) in comparison with decidua (DC)), were identified by differential expression gene analysis. The transcription factors enriched in these models were then identified by newly developed tools BART [8] tool, which provides functional interpretations to differential gene expression analysis. Figure 1 shows the experiment and analysis process of this study. Table 5 Summary of datasets analyzed in this study.

**Figure 1.**
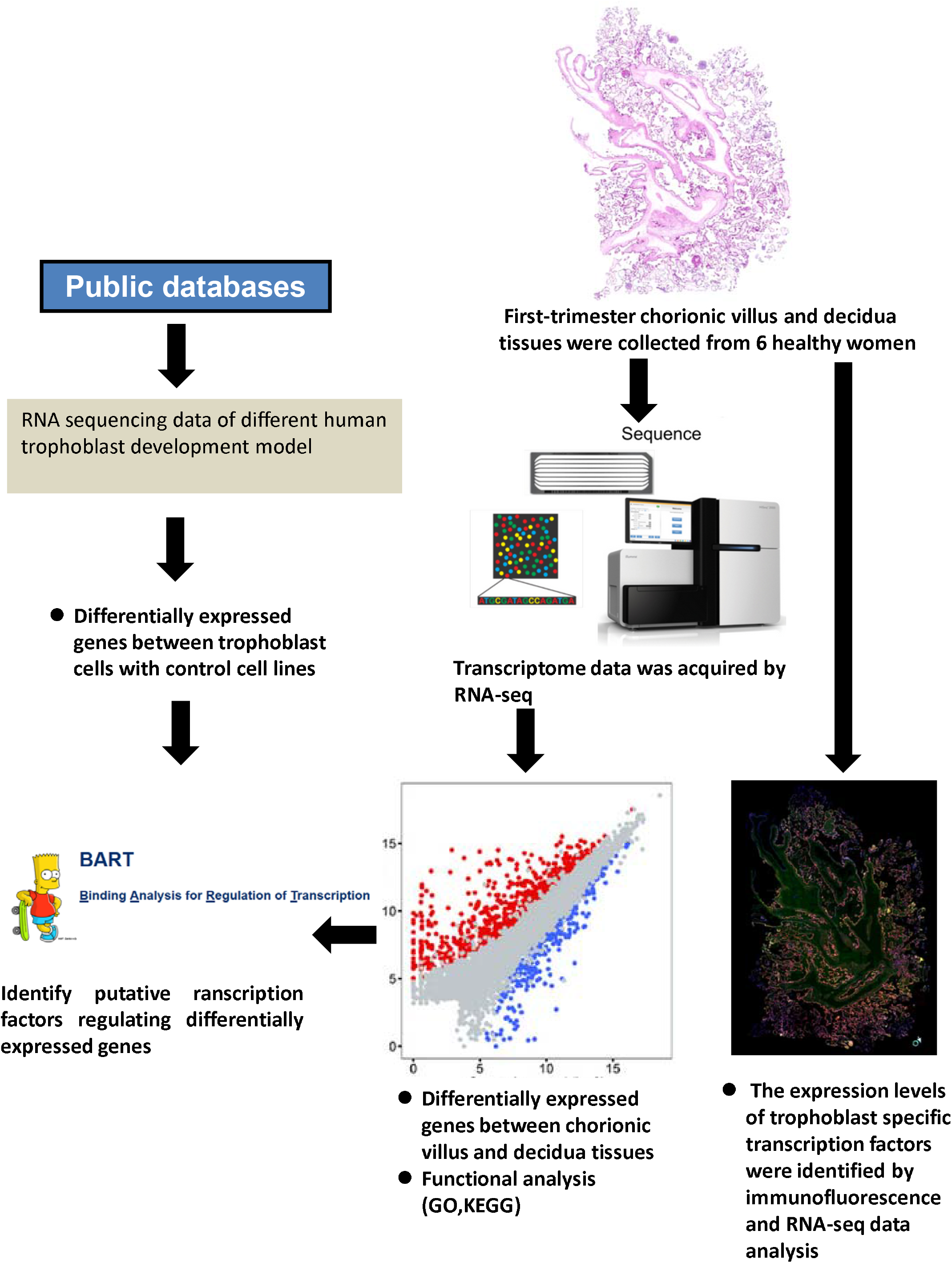
The experiment and analysis workflow of this study.

## Materials and Method

### Collection protocol

The first-trimester placenta (six to nine-week of gestation from 6 healthy women, confirmed by the embryo size under detection of ultrasound) was collected in the Second Affiliated Hospital of Zhengzhou University. After being separated ex-vivo, the placenta was immediately washed in the ice saline and divided into several parts by a scalpel blade on ice, and then were transformed into cryogenic vials which were filled with 1 ml RNA store beforehand. When disposed of properly, the samples were put into 4°C refrigerator for 24h to let the RNA store immerse them. After 24h the RNA store was abandoned and the samples were sopped up using sterile absorbing paper, then the samples were separately collected into new cryogenic vials and were store into −80°C refrigerator for further investigations.

### HE (Hematoxylin & Eosin) staining

Samples collected before were fixed in formalin, embedded in paraffin and sliced up to 4 μm sections, and then were deparaffinized and rehydrated. The deparaffin and rehydration protocols are xylene I for 5 min, xylene II for 5 min, 100% ethanol for 2min, 95% ethanol for 1min, 80% ethanol for 1min, 75% ethanol for 1min, and finally distilled water for 2min. After the process above, the sections were stained in hematoxylin for 5 min and rinsed with tap water, then differentiated in hydrochloric acid and ethanol for the 30s respectively. At last, the sections were soaked into tap water for 5min and sealed in neutral resins with cover glass, then were observed under an ordinary optical microscope, 10 pictures were randomized obtained per section.

### Immunohistochemistry

Paraffin-embedding, deparaffinized, and the rehydrating process is the same as HE staining. After these steps, the sections were subjected to antigen retrieval for 3min in a medical pressure cooker with citration solution (pH=6), subsequently treated with endogenous catalase blocker and horse serum to eliminate the interference of endogenous catalase and nonspecific staining. The sections were then incubated with primary antibody. Next day the sections were washed in PBS and then incubated in the secondary antibody (1:200, Abbkine) for 1 hour in room temperature, and after DAPI staining, the sections were observed under the fluorescence microscope, pictures were randomized obtained per section.

### RNA-seq experiment

Total RNA was extracted with Trizol (Tiangen, Beijing) and assessed with Agilent 2100 BioAnalyzer (Agilent Technologies, Santa Clara, CA, USA) and Qubit Fluorometer (Invitrogen). Total RNA samples that meet the following requirements were used in subsequent experiments: RNA integrity number (RIN) > 7.0 and a 28S:18S ratio > 1.8. RNA-seq libraries were generated and sequenced by CapitalBio Technology (Beijing, China). The triplicate samples of all assays were constructed an independent library, and do the following sequencing and analysis. The NEB Next Ultra RNA Library Prep Kit for Illumina (NEB) was used to construct the libraries for sequencing. NEB Next Poly(A) mRNA Magnetic Isolation Module (NEB) kit was used to enrich the poly(A) tailed mRNA molecules from 1 μg total RNA. The mRNA was fragmented into ∼200 base pair pieces. The first-strand cDNA was synthesized from the mRNA fragments reverse transcriptase and random hexamer primers, and then the second-strand cDNA was synthesized using DNA polymerase I and RNaseH. The end of the cDNA fragment was subjected to an end repair process that included the addition of a single “A” base, followed by ligation of the adapters. Products were purified and enriched by polymerase chain reaction (PCR) to amplify the library DNA. The final libraries were quantified using KAPA Library Quantification kit (KAPA Biosystems, South Africa) and an Agilent 2100 Bioanalyzer. After quantitative reverse transcription-polymerase chain reaction (RT-qPCR) validation, libraries were subjected to paired-end sequencing with pair-end 150-base pair reading length on an Illumina NovaSeq 6000.

### RNA-seq data analysis

Transcript abundance was quantified using Kallisto (Bray et al., 2016) and gene fold changes were generated by comparing gene expression levels between two groups using the limma R package (Ritchie et al., 2015). Figure S3 shows library size analysis results. P-value or q-value was used to conduct a significance analysis. Parameters for classifying significantly DEGs are >2-fold differences (|log2FC|>1, FC: the fold change of expressions) in the transcript abundance and q < 0.05. Gene fold changes were transformed using log2 and displayed on the x-axis; P-values were corrected using the Benjamini-Hochberg method, transformed using –log10, and displayed on the y-axis. The functional enrichment analysis was performed using g: Profiler (version e99_eg46_p14_f929183) with g: SCS multiple testing correction methods applying a significance threshold of 0.05 (Raudvere et al., 2019). Hierarchical clustering of arbitrary types of objects from a matrix of distances and shows a corresponding dendrogram based on Orange (Demšar et al., 2013). The parameter setting could be found in Figure S8.

### 2D mapping

Orange [11], which provided a wrapper for scikit-learn algorithms [12], was used for batch effect remove, filtering (by cells and genes), scaling, normalization, clustering, dimensionality reduction, clustering and visualize cell clusters using, t-SNE, and PCA.

### Identify putative transcription factors regulating differentially expressed genes

We used the transcription factor prediction tool BART ^12^. BART was run with all default settings, and the provided transcription factor databases. BART was run with all default settings, and the provided transcription factor databases.

## Results

### The expression level of trophoblast specific transcription factors in Chorionic villus

Chorionic villus exhibits typical chorionic villus tissue morphology according to hematoxylin and eosin stain (H&E tissue sections) (Figure 2A). We have previously identified specific transcription factors (including *JUN, FOS, TFAP2A, TFAP2C, TEAD1, TEAD3, TEAD4, GATA2, GATA3*) in trophoblast cells based on DNas-seq derived from ENCODE project [14]. The results showed that *JUN, FOS, TFAP2A, TFAP2C, TEAD4, GATA2, GATA3* were most strongly expressed in chorionic villus (Figure 2B). *TEAD1, TEAD3* can be detected, though at lower levels than other genes. As a control, the well-studied chorionic villus surface marker *HLA-G* and *KRT7* were also expressed in the chorionic villus (Figure S1A, B).

**Figure 2.**
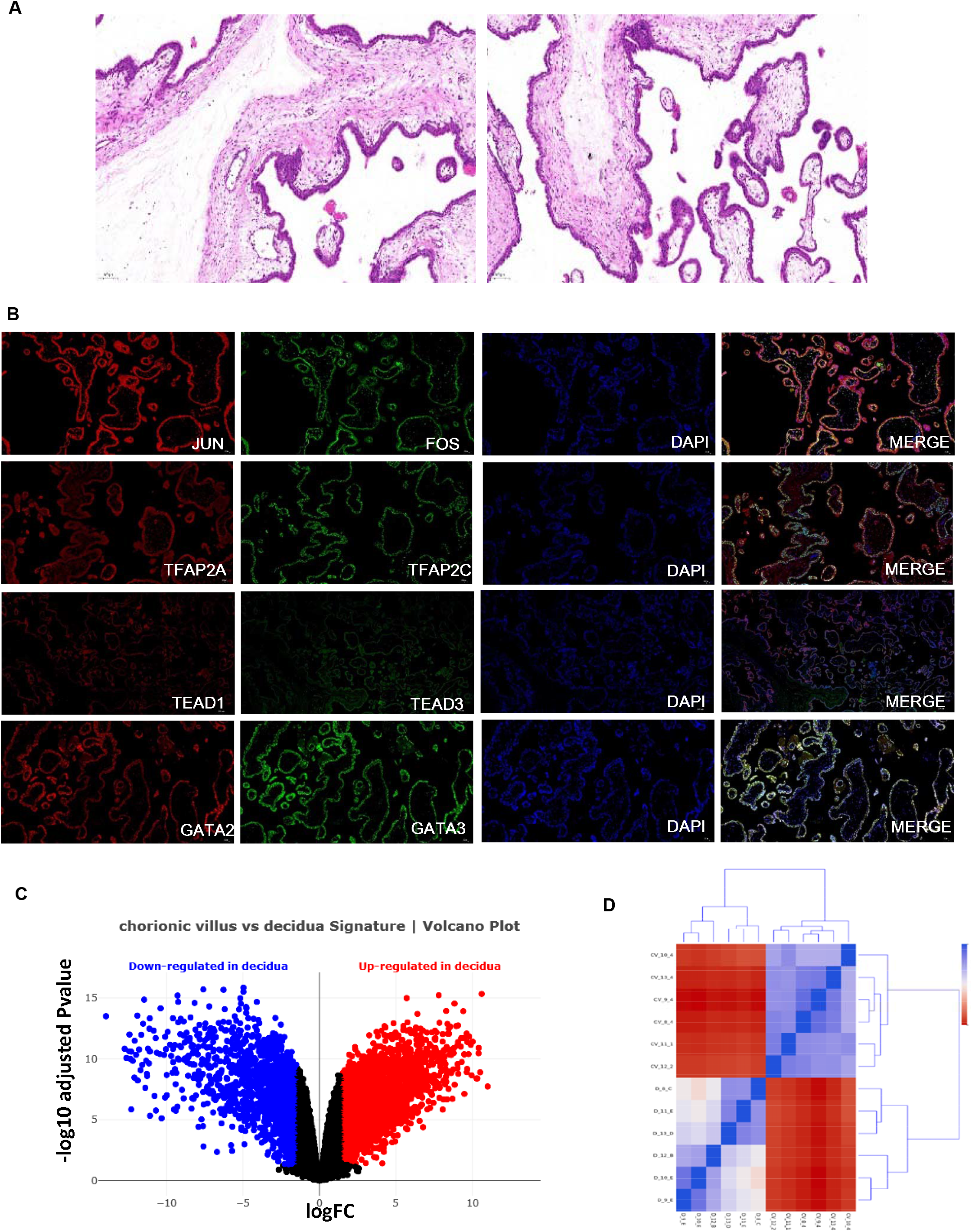
(A) Tissue section, hematoxylin staining in chorionic villus; (B) Immunofluorescence staining of the trophoblastic specific transcription factor in chorionic villus; (C) Scatter plots of differentially expressed genes between chorionic villus and decidua. Every point in the plot represents a gene. Red points indicate significantly up-regulated genes, and blue points indicate down-regulated genes; (D) Hierarchical clustering and heatmap displaying differentially expressed genes between chorionic villus and decidua. Every row of the heatmap represents a gene, every column represents a sample, and every cell displays normalized gene expression values;

### RNA-Sequencing of first-trimester chorionic villus and decidua

Transcriptome data of chorionic villus and decidua from six first-trimester donors were obtained by high-throughput sequencing. Table 1 listed the top 10 highly expressed genes in six human chorionic villus tissues. Among them, human chorionic gonadotropin (hCG) family genes including *CGA, CGB5, CGB8, CGB3*, and pregnancy-specific glycoproteins (PSGs) [16] family genes including *PSG3, PSG1* are the most abundantly expressed genes in chorionic villus samples (Table S1, Figure S2A). Table 2 listed the top 10 highly expressed genes in corresponding decidua tissues.

**Table 1.**
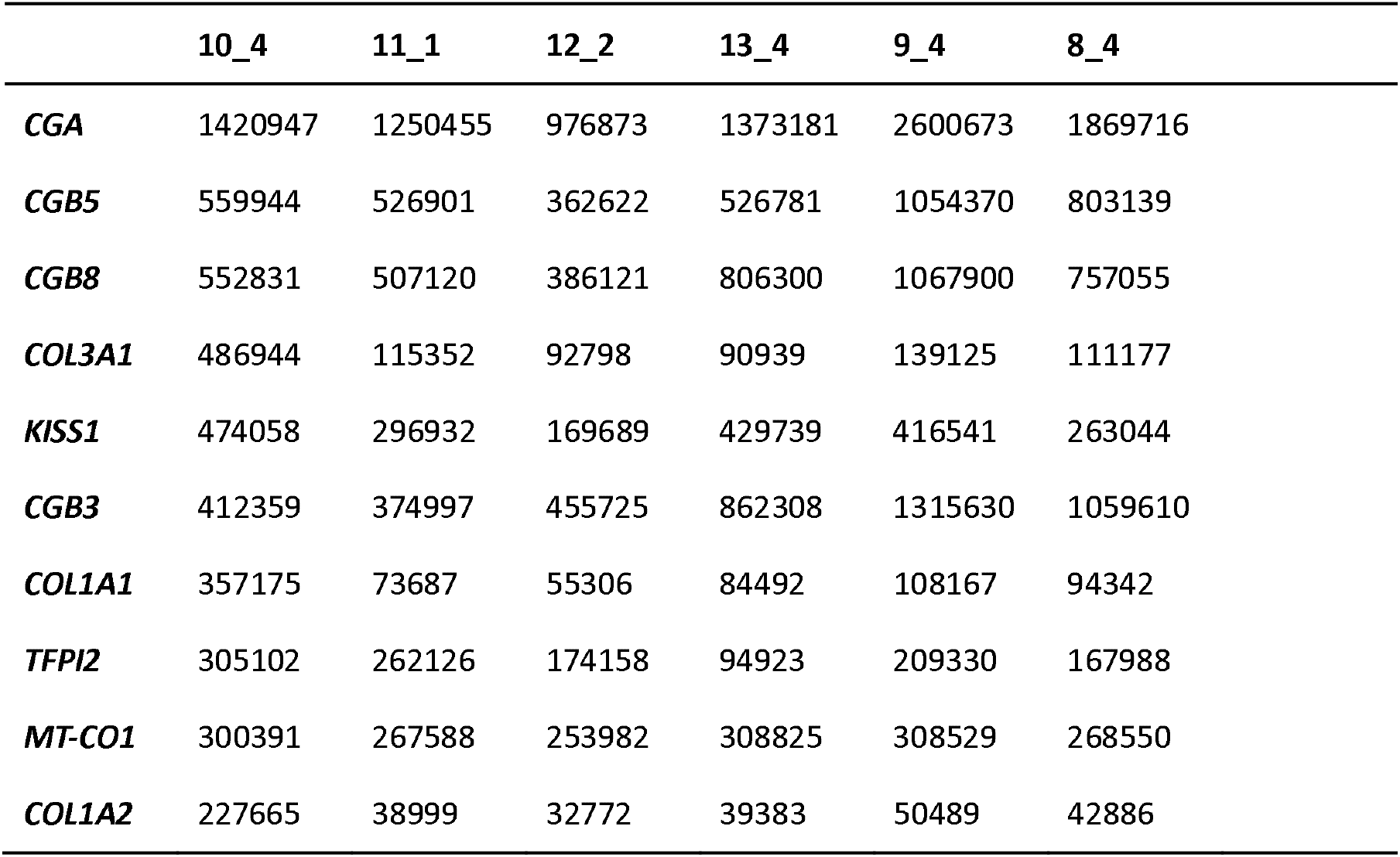
Top 10 genes that are highly expressed in the expression profile of villus tissue. Rows represent genes, columns represent samples, and values show the number of mapped reads.

|  | 10_4 | 11_1 | 12_2 | 13_4 | 9_4 | 8_4 |
| --- | --- | --- | --- | --- | --- | --- |
| ***CGA*** | 1420947 | 1250455 | 976873 | 1373181 | 2600673 | 1869716 |
| ***CGB5*** | 559944 | 526901 | 362622 | 526781 | 1054370 | 803139 |
| ***CGB8*** | 552831 | 507120 | 386121 | 806300 | 1067900 | 757055 |
| ***COL3A1*** | 486944 | 115352 | 92798 | 90939 | 139125 | 111177 |
| ***KISS1*** | 474058 | 296932 | 169689 | 429739 | 416541 | 263044 |
| ***CGB3*** | 412359 | 374997 | 455725 | 862308 | 1315630 | 1059610 |
| ***COL1A1*** | 357175 | 73687 | 55306 | 84492 | 108167 | 94342 |
| ***TFPI2*** | 305102 | 262126 | 174158 | 94923 | 209330 | 167988 |
| ***MT-CO1*** | 300391 | 267588 | 253982 | 308825 | 308529 | 268550 |
| ***COL1A2*** | 227665 | 38999 | 32772 | 39383 | 50489 | 42886 |

**Table 2.** Top 10 genes that are highly expressed in the expression profile of decidua. Rows represent genes, columns represent samples, and values show the number of mapped reads.

|  | 10_E | 11_E | 12_B | 13_D | 8_C | 9_E |
| --- | --- | --- | --- | --- | --- | --- |
| ***PAEP*** | 707159 | 225495 | 553927 | 332730 | 317333 | 716464 |
| ***MT-CO1*** | 402678 | 380707 | 397730 | 315579 | 310992 | 375237 |
| ***EEF1A1*** | 274212 | 194122 | 314425 | 233226 | 203655 | 228912 |
| ***FBLN1*** | 148393 | 46328 | 95584 | 46766 | 46616 | 108020 |
| ***MT-ND4*** | 127854 | 100057 | 153752 | 90144 | 108484 | 126805 |
| ***MT-CO3*** | 118360 | 84003 | 113272 | 77937 | 63105 | 121600 |
| ***MT-ND5*** | 112658 | 86210 | 65977 | 83994 | 96545 | 104019 |
| ***MT-CYB*** | 108860 | 74152 | 99149 | 66198 | 63807 | 81149 |
| ***GPX3*** | 103827 | 113570 | 280574 | 105654 | 71261 | 78752 |
| ***IGFBP5*** | 88947 | 49685 | 30649 | 33556 | 25273 | 108734 |

Then, we identify the sex of the cell line based on the recently determined gender-specific transcript markers, *RPS4Y1, EIF1AY, DDX3Y, KDM5D* (Staedtler et al., 2013). The results indicate that 10_4, 11_1, 12_2 is derived from males, and 13_4, 9_4 is derived from the female (Figure S2B). We then compared the differentially expressed genes between the chorionic villus (13_4, 9_4, 10_4, 11_1, 12_2) and decidua (10_E,11_E,12_B,13_D,8_C,9_E) groups. Volcano plots are used to display the differential expressions of genes in each group (Figure 2C). Each point in the scatter plot represents a gene; the axes display the significance versus fold-change estimated by the differential expression analysis. Our analysis identified that 1315 genes were up-regulated in chorionic villus compared with decidua and 2272 genes were up-regulated in decidua compared with chorionic villus (log2FoldChange, p<0.05) (Table S3). The 10 top genes (logFC>2, p<0.05) significantly upregulated in chorionic villus compared with decidua are shown in Table 3. The top 10 genes (logFC>2, p<0.05) significantly up-regulated in decidua compared with chorionic villus are shown in Table 4. Kisspeptin and its receptors, highly expressed in chorionic villus, play an essential role in the establishment of the maternal-fetal dialogue.

**Table 3.** The 10 top genes significantly upregulated in chorionic villus compared with decidua (logFC>2, p<0.05). Every row of the table represents a gene; the columns display the estimated measures of differential expression.

| gene_symbol | logFC | AveExpr | t | P.Value | adj.P.Val |
| --- | --- | --- | --- | --- | --- |
| ***DLK1*** | -13.9637 | 2.418015 | -32.4705 | 3.16E-14 | 1.83E-11 |
| ***KISS1*** | -12.7453 | 7.639202 | -20.4116 | 1.49E-11 | 8.85E-10 |
| ***PSG8*** | -12.6297 | 2.410962 | -17.86 | 8.54E-11 | 3.14E-09 |
| ***CGA*** | -12.4301 | 10.01645 | -19.6388 | 2.47E-11 | 1.28E-09 |
| ***XAGE3*** | -12.4281 | 0.951458 | -25.0237 | 1.01E-12 | 1.46E-10 |
| ***CGB5*** | -12.3711 | 8.724329 | -17.6354 | 1.01E-10 | 3.50E-09 |
| ***CGB2*** | -12.3182 | 3.340725 | -8.13134 | 1.43E-06 | 6.77E-06 |
| ***DUSP9*** | -12.311 | 2.240281 | -19.5565 | 2.61E-11 | 1.30E-09 |
| ***CGB3*** | -12.2089 | 8.946367 | -17.1212 | 1.48E-10 | 4.74E-09 |
| ***CGB7*** | -12.1083 | 3.189653 | -13.386 | 3.46E-09 | 5.30E-08 |

**Table 4.**
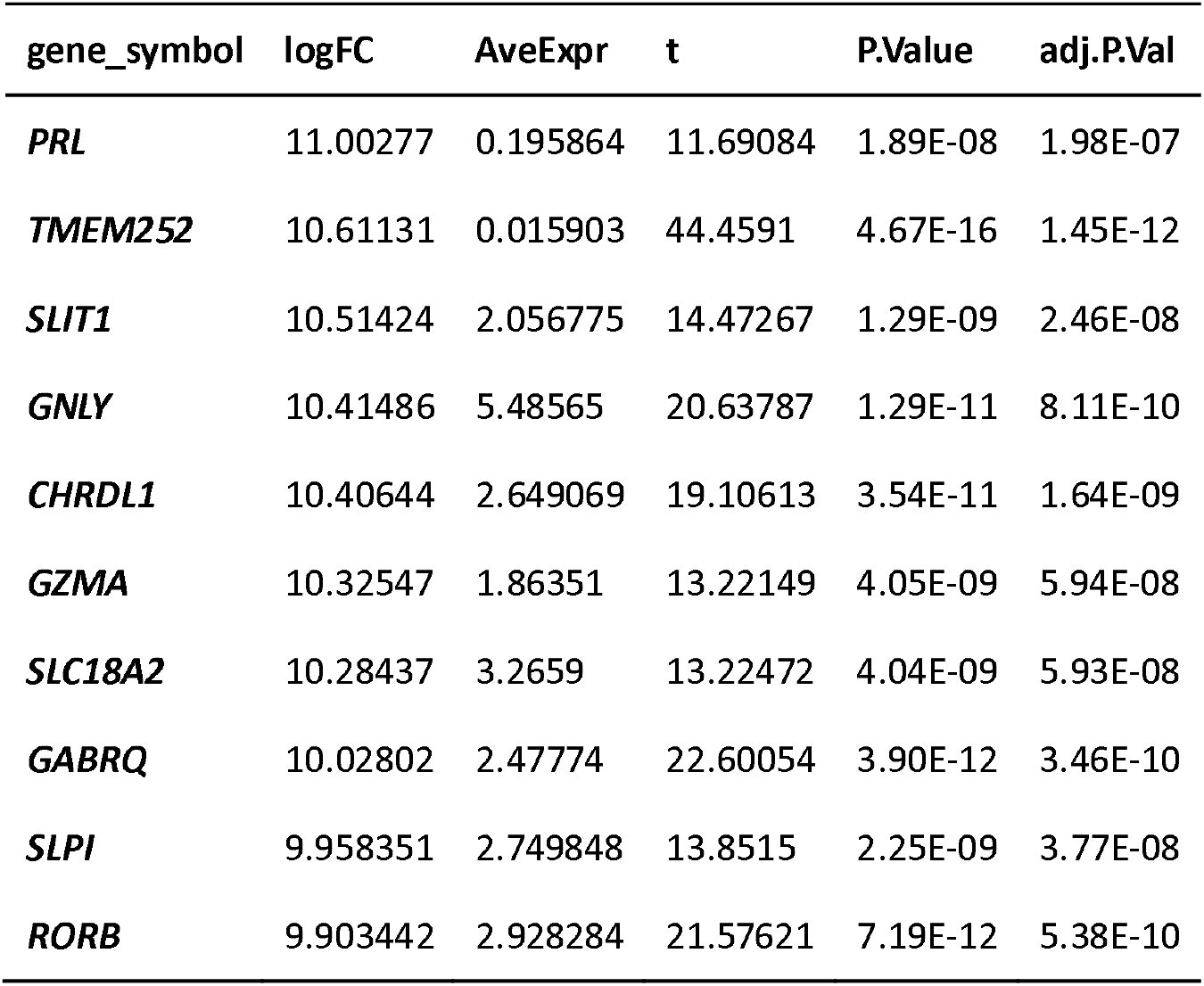
The 10 top genes significantly upregulated in chorionic villus compared with decidua (logFC>2, p<0.05). Every row of the table represents a gene; the columns display the estimated measures of differential expression.

Next, we analyzed the expression of trophoblastic cell-specific transcription factors (Knott and Paul, 2014) and putative surface marker genes in chorionic villus (Figure 2D, 2E). Our analysis was based on the system of stem cell differentiation into trophoblast cells (ESCd1 (Yabe et al., 2016), ESCd2 (Krendl et al., 2017)), immortalized trophoblast cells (HTR8/SVneo (Lee et al., 2016), Bewo (Renaud et al., 2015), Jeg3 (Ferreira et al., 2016)), and the chorionic villus collected in this study. For the corresponding gene expression matrix, please refer to Table S2. Interestingly, genes from chorionic gonadotropin (hCG) family genes are highly expressed in both chorionic villus and cell lines. While, pregnancy-specific glycoproteins (PSGs) family genes are highly expressed only in chorionic villus, and at a low expression level in vitro cell. Trophoblast-specific transcription factors are highly expressed in chorionic villus and showed heterogeneous expression level in immortalized cell lines and stem cell differentiation systems. Immunofluorescence results confirmed that *TFAP2C* was detectable in immortalized trophoblastic cell lines JEG3 in vitro (Figure S1C).

Enrichment analysis revealed enriched GO terms in the top 500 upregulated genes in chorionic villus (Figure S4) including “reproductive process”, “multicellular organismal process”, “hormone activity”, “animal organ morphogenesis”, “female pregnancy”, “embryonic morphogenesis”, “placenta development”, “cis-regulatory region binding”[17] et al. Not surprisingly, enriched GO terms of upregulated genes in the decidua, the primary place for the establishment and communication of pregnancy immune microenvironment, include many immune-related terms such as “regulation of immune system process”, “positive regulation of immune system process” and “immune response” et al (Figure S5).

KEGG pathway analysis of the top 500 upregulated genes in chorionic villus revealed previously identified trophoblast related pathways “PPAR signaling pathway” (Barak et al., 2008), (Figure S4). KEGG analysis of the top 500 upregulated genes in decidua shows that immune-related pathways approximately occupying the top ten pathways, including” Antigen processing and presentation”, “Cytokine-cytokine receptor interaction”, and “Th17 cell differentiation” (Figure S5). More detailed analysis results on GO and KEGG enrichment analysis could be found in Supplementary file 1.

### PCA and t-SNE function similarly in identifying a small number of samples

PCA and t-SNE analysis was employed to identify global patterns of the gene expression profile of six chorionic villus and corresponding decidua tissues. The results show an overall similarity result generated by PCA and the t-SNE map (Figure 3A). Both methods correctly clustered 12 samples into two types, namely chorionic villus and decidua. Nevertheless, t-SNE can distinguish these two groups more significantly into different tissue types, and the same type of tissues is more closely clustered together (Figure 3B).

**Figure 3.**
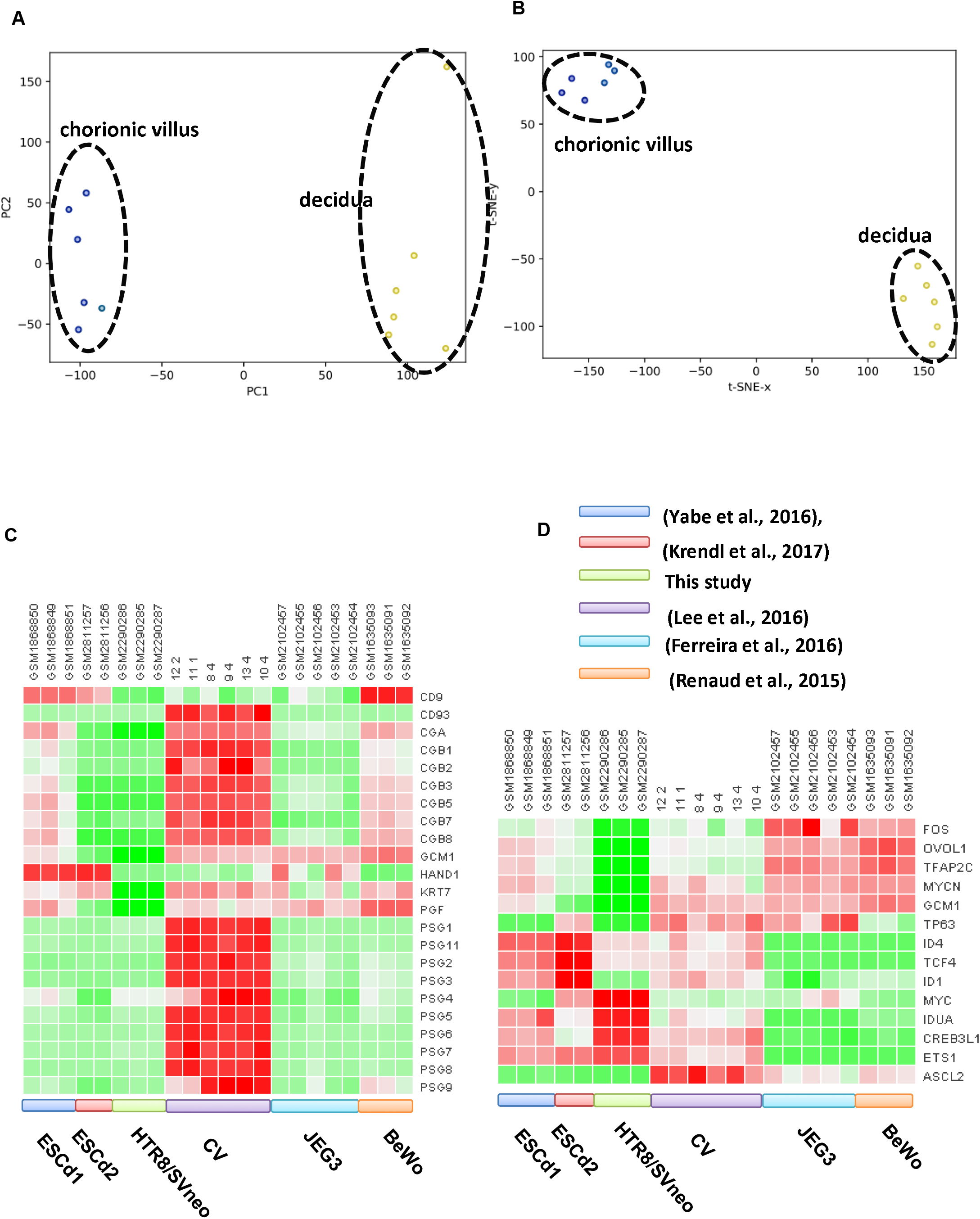
(A) Projection of 12 samples (including chorionic villus and decidua) in a 2D-map using PCA: Each point represents a sample which is colored according to the (sub)tissue label; (B) Projection of 12 samples (including chorionic villus and decidua) in a 2D-map using t-SNE (Perplexity=11; metric was set as Euclidean); (C) Expression of pregnancy-related surface marker genes and transcription factors (D) in different cell models of chorionic villus (CV) based on RNA-seq data analysis.

### Comparison of the transcriptome of different trophoblast cell models

First, genes up-regulated in different trophoblastic cell models were obtained by differential expression gene analysis, including hESC line cells after BMP4 treatment (TB) comparison with H1^5^ and trophectoderm (TE) in comparison with pluripotent epiblast (EPI) cells ^6^, chorionic villus (CV) in comparison with decidua (DC)) (Table S4, 5). We chose the top 200 different expression from each group to predict the transcriptional regulators of trophoblastic cells using a newly developed functional transcriptional regulatory predictive tool [8]. Different cell models showed different enrichment of transcription factors (Figure 4A). *GATA3, TRIM28*, and *ESR1* is the most enriched transcription factor in TB, CV and TE respectively (Figure 4B). All the trophoblastic cells from three different sources were enriched *GATA3, GRHL2, NR3C1, EP300, TP63, BANF1, TEAD4, ESR1, TFAP2A, TFAP2C, TP53, YAP1, H2AFX*. These transcription factors can be used as components of the core transcriptional regulatory network of trophoblast cells. In general, CV and TB were relatively close, with more overlapping transcription factors, while TE had less overlapping transcription factors. The intersection and complement of these data sets are listed in Table 6.

**Table 5.** Summary of datasets analyzed in this study.

| Cell lines | GEO<br>number | Description | Reference |
| --- | --- | --- | --- |
| HTR8/SVneo | GSE85995 | Function and hormonal regulation of GATA3 in human first-trimester placentation | [18] |
| JEG3 | GSE79779 | Transcriptional profiling of JEG3 cells with HLA-G ablation via deletion of Enhancer L | [19] |
| hESC line cells after BMP4 treatment | GSE73017 | Comparison of syncytiotrophoblast differentiated from human pluripotent stem cells and that derived from primary cytotrophoblast recovered from term placentae | [20] |
| hESC line cells after BMP4 treatment | SRP000941 | H1 cells were differentiated to trophoblast-like cells | [6] |
| Trophectoderm | GSE66507 | Single-Cell RNA-seq Defines the Three Cell Lineages of the Human Blastocyst | [7] |
| Chorionic villus | ERX3472341 | chorionic villus in early-gestation stage | This study |
| Decidua | ERX3472299 | decidua in early-gestation stage | This study |

**Figure 4.**
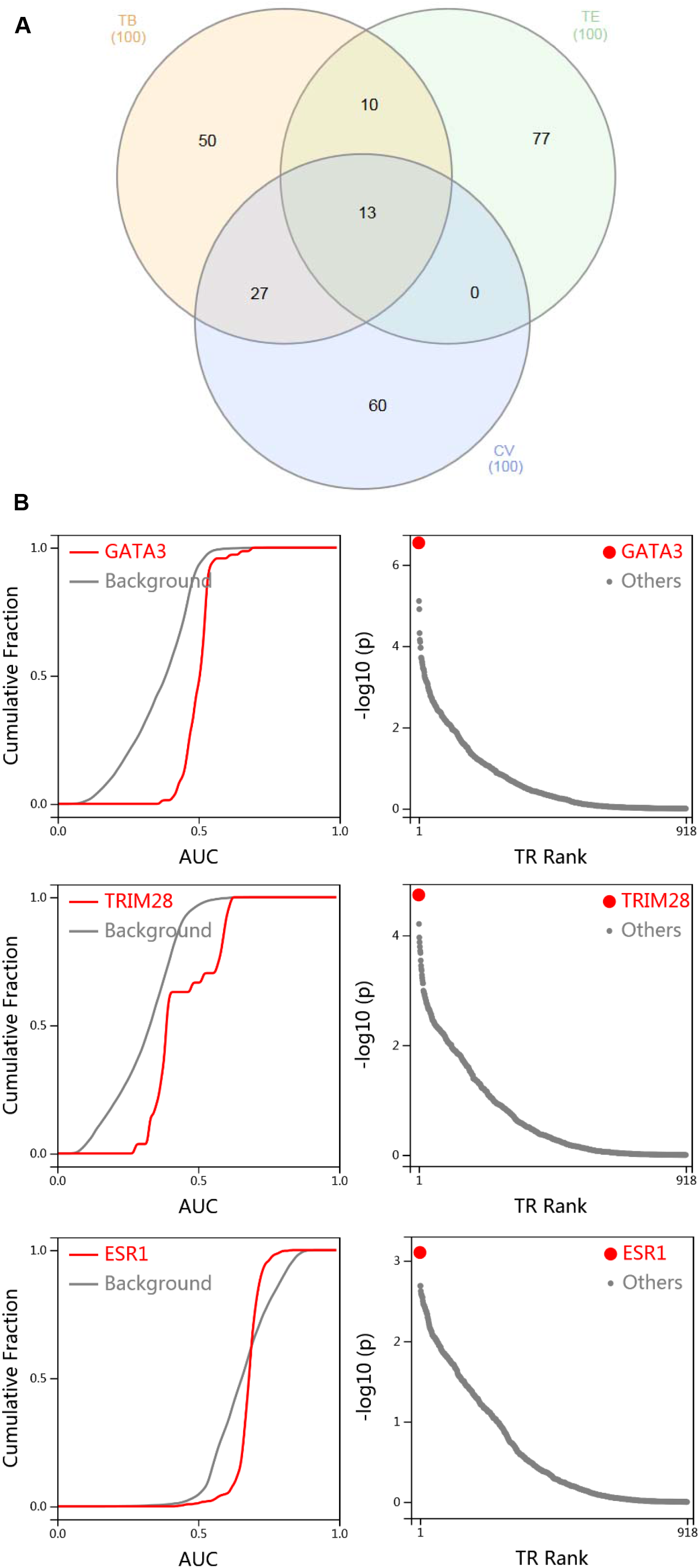
(A)Venn diagram of top 100 enriched transcription factors in different trophoblastic cell models by BART; (B) Top enriched transcription factors in different trophoblastic cell models (TB, CV, and TE respectively) by BART; Area under the ROC curve (AUC) is calculated for each dataset; AUC are grouped by the factor, and Wilcoxon test is performed for each factor compared with all datasets as background. cumulative distributions show significantly higher AUC for TF.

**Table 6.**
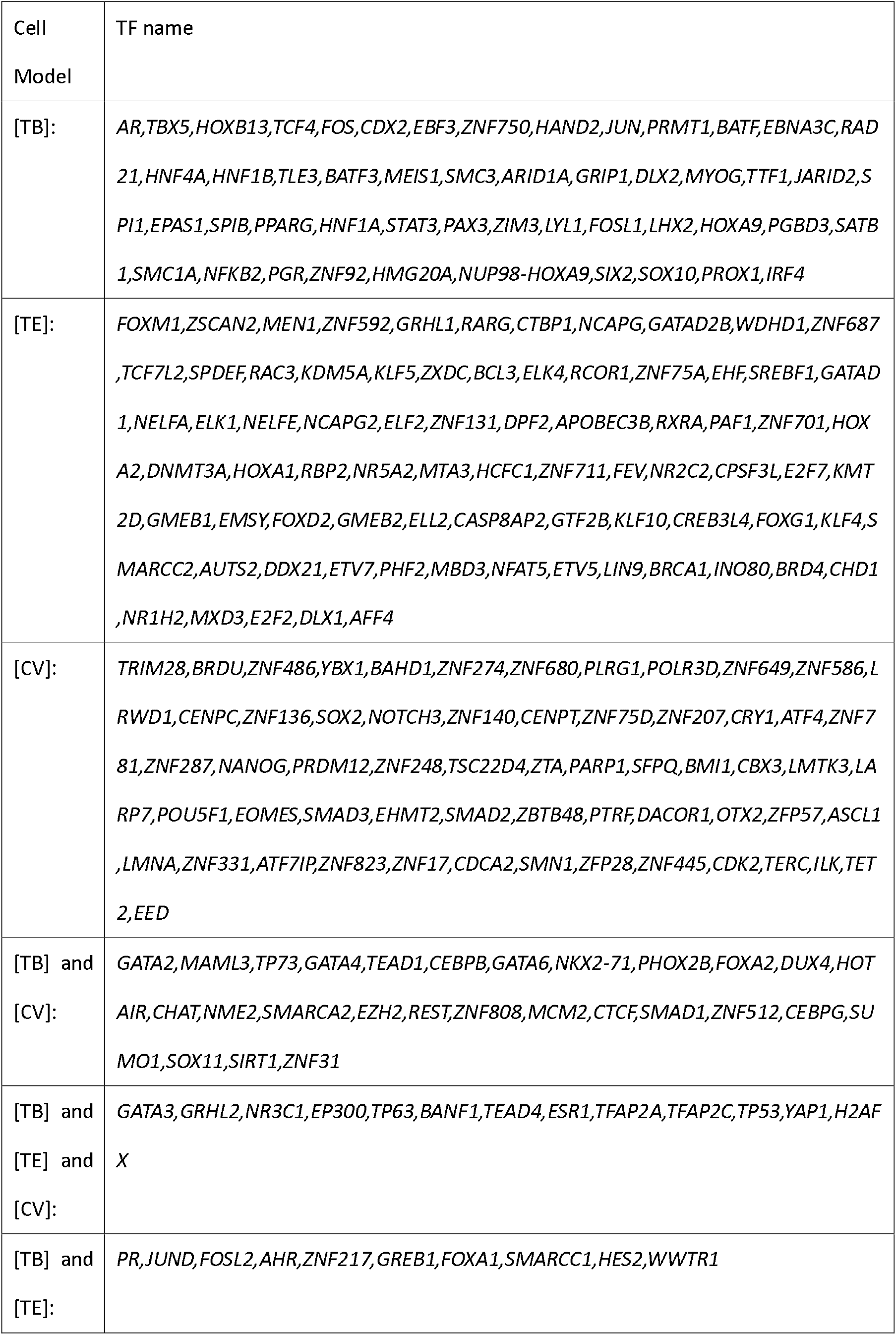
TFs enriched in different trophoblast cell models (TB: hESC line cells after BMP4 treatment, TE: Trophectoderm, CV: Chorionic villus).

## Discussion

In this study, we collected six human chorionic villus and decidua tissue at an early stage of pregnancy and performed transcriptome sequencing based on high-throughput sequencing technology. Although there are public RNA-seq data available for chorionic villus and decidua tissue, this is the first time that the RNA-seq data were obtained from the chorionic villus and decidua which derived from the same patient.

We obtained trophoblastic cell-specific transcription factors based on chromatin accessibility data in previous studies [14]. In this study, we are based on a recently developed transcription factor prediction tool BART on transcriptome data obtained from the same article and process [6]. Based on these two data sets, many transcription factors have been successfully predicted (for example, *GATA3, GATA2, TEAD1, JUND, TEAD4, FOS, TFAP2A, JUN, TFAP2C* et al). The analysis indicates the reliability of the analysis results of this newly developed tool. The experiment and cost of transcriptome data acquisition are significantly lower than Dnas-seq and ChIP-seq. Therefore, this tool provides a technical guarantee for convenient transcriptional regulation analysis in the future.

We identified genes that were relatively highly expressed in trophoblast cells in different trophoblast cell models. At the same time, we identified transcription factors that these genes might be regulated. This analysis is of certain significance for further exploration of the development of placenta and the occurrence of pregnancy-related diseases in the future.

## Acknowledgments

All authors contributed to the study conception and design. LYJ conceived the project and completed the core program. LYJ performed the computational analysis. ZY and YLG performed the wet experiment. HGM, RJL, and MJ provide the necessary software and hardware foundation for this research. LYJ wrote the manuscript. All authors analyzed and discussed the results.

We give thanks to Qunying Wei of Department of Obstetrics and Gynecology, the Second Affiliated Hospital of Zhengzhou University for technical support. We also give thanks to Wuhan ServiceBio technology co.LTD for the assistance on the immunohistochemical experiment. Sequencing results from this study have been assigned to European Nucleotide Archive with accession ERX3472341 for chorionic villus and ERX3472299 for decidua.

## Supplementary table

Table S1 gene expression matrix of trophoblastic cell-specific transcription factors and putative surface marker genes in chorionic villus;

Table S2 Gene expression profiles of 6 Villus and decidua derived by this study;

Table S3 differentially expressed genes between chorionic villus vs decidua;

Table S4 differentially expressed genes between TB and H1;

Table S5 differentially expressed genes between TE and EPI.

## Supplementary file

Supplementary file 1: GO and KEGG enrichment analysis of differentially expressed genes between chorionic villus and decidua;

## Supplementary figure

Figure S1 (A) Immunostaining of HLA-G in chorionic villus; (B) Immunofluorescence of trophoblast-specific surface markers KRT7 and HLA-G in Chorionic villus; (C) Immunofluorescence results of TFAP2C in immortalized trophoblastic cell lines (JEG3) in vitro.

Figure S2 (A) Track of the gene expression signal (intensity on Y-axis) at human chorionic gonadotropin (hCG) family and pregnancy-specific glycoproteins (PSGs) in chorionic villus tissues in the human genome; (B) Expression level of tissues-specifically expressed genes, RPS4Y1, EIF1AY, DDX3Y and KDM5D in different chorionic villus tissues;

Figure S3 Library Size Analysis results. The figure contains an interactive bar chart that displays the total number of reads mapped to each RNA-seq sample in the dataset.

