## Supplementary material for "Comparison analysis on transcriptomic of different human trophoblast development model": Supplementary figure.pptx

### Slide 1
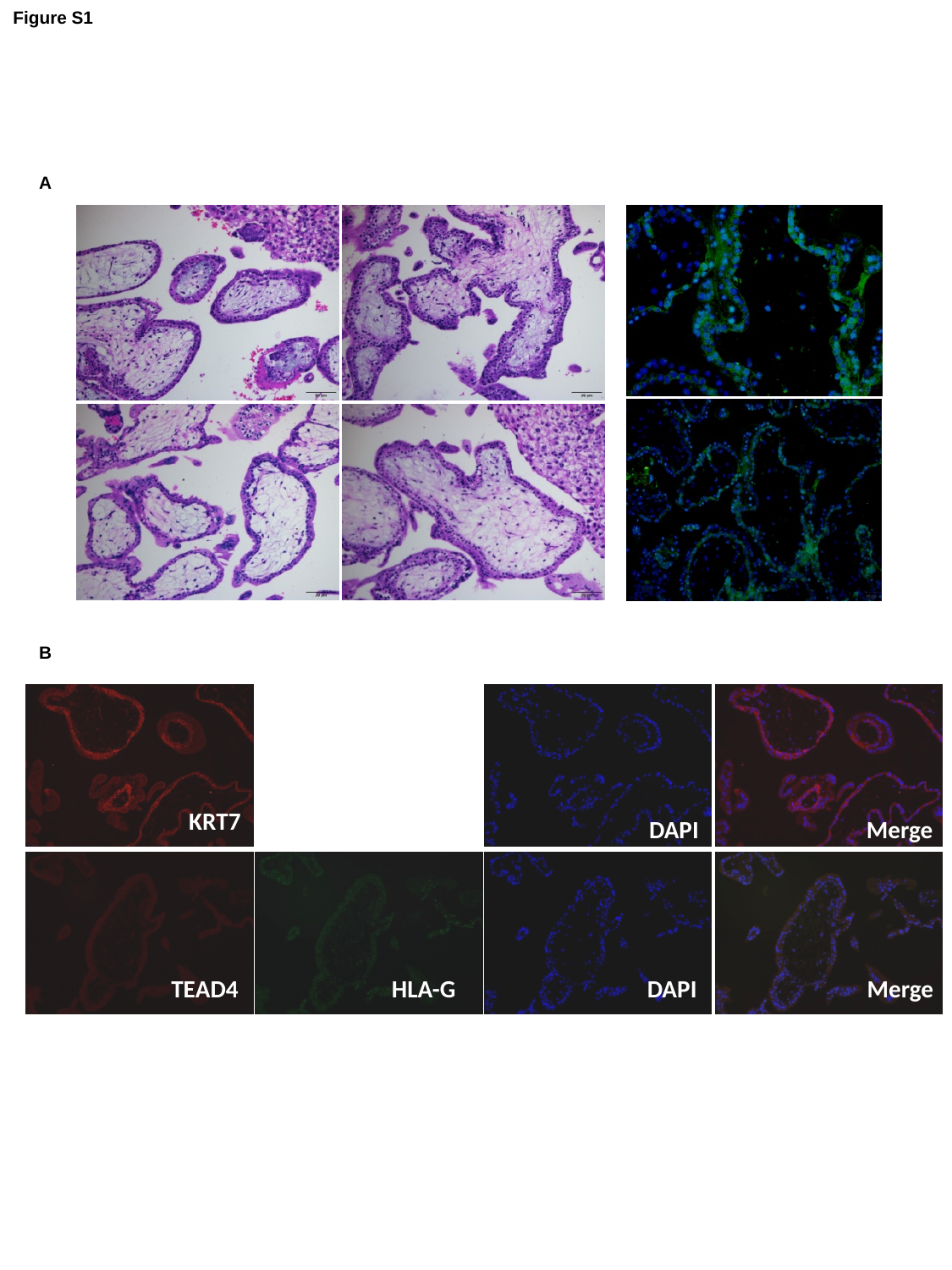

Figure S1
A
B
KRT7
DAPI
Merge
TEAD4
HLA-G
DAPI
Merge

### Slide 2
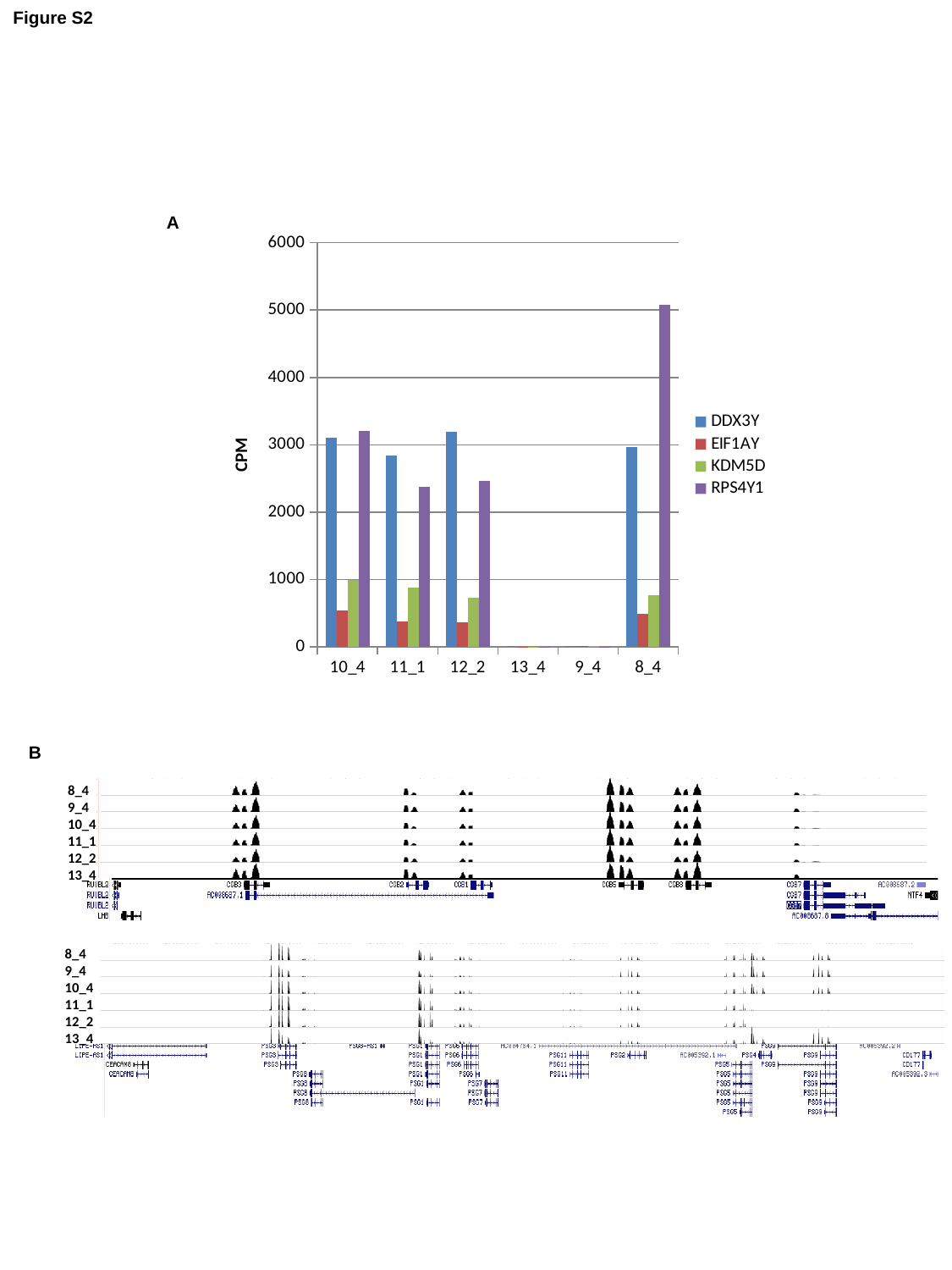

Figure S2
A
#### Chart
| Category | DDX3Y | EIF1AY | KDM5D | RPS4Y1 |
|---|---|---|---|---|
| 10_4 | 3109.0 | 534.0 | 988.0 | 3210.0 |
| 11_1 | 2841.0 | 378.0 | 878.0 | 2374.0 |
| 12_2 | 3195.0 | 357.0 | 723.0 | 2465.0 |
| 13_4 | 5.0 | 3.0 | 4.0 | 3.0 |
| 9_4 | 5.0 | 13.0 | 13.0 | 3.0 |
| 8_4 | 2970.0 | 488.0 | 771.0 | 5081.0 |B
8_4
9_4
10_4
11_1
12_2
13_4
8_4
9_4
10_4
11_1
12_2
13_4

### Slide 3
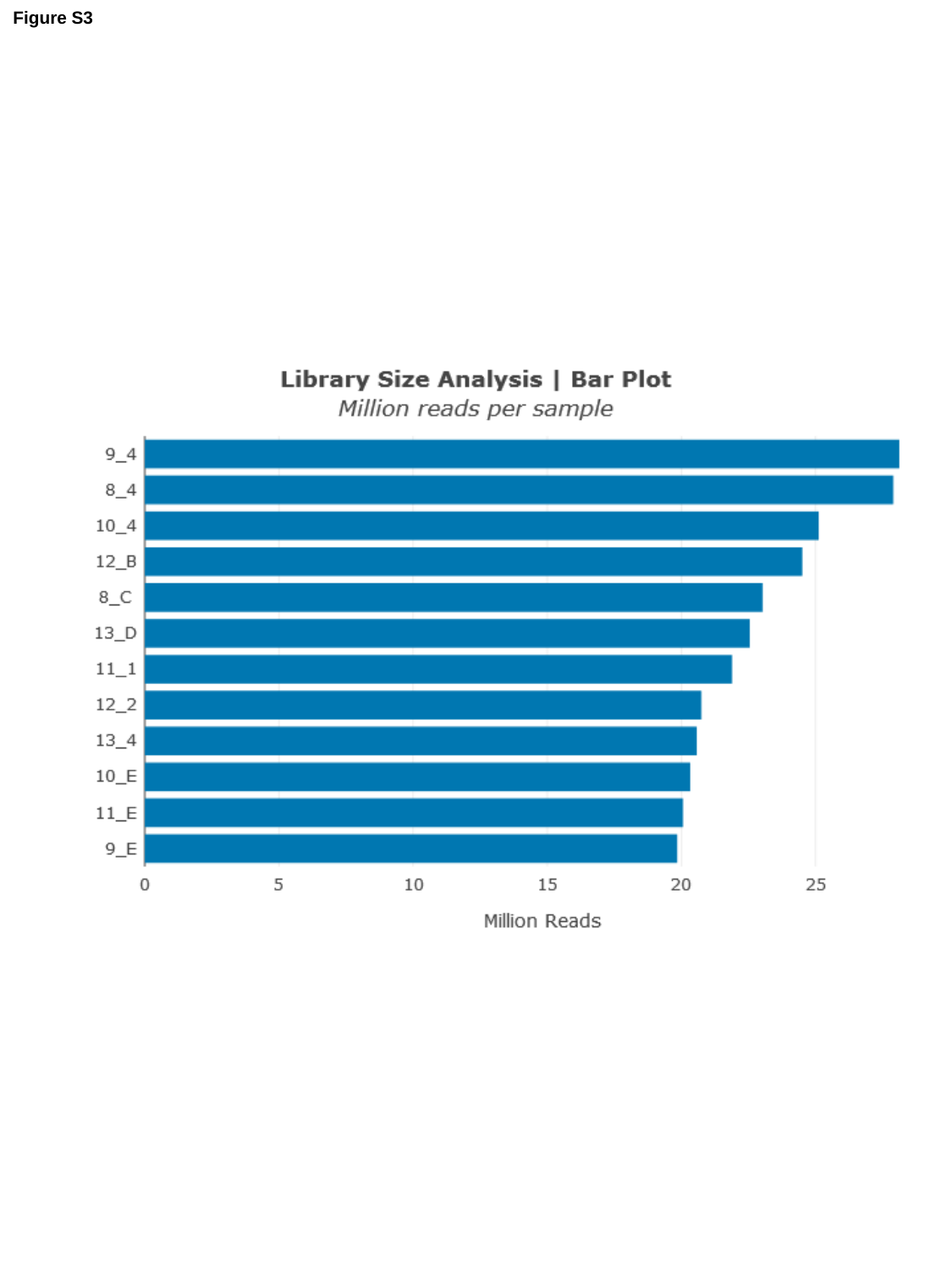

Figure S3

### Slide 4
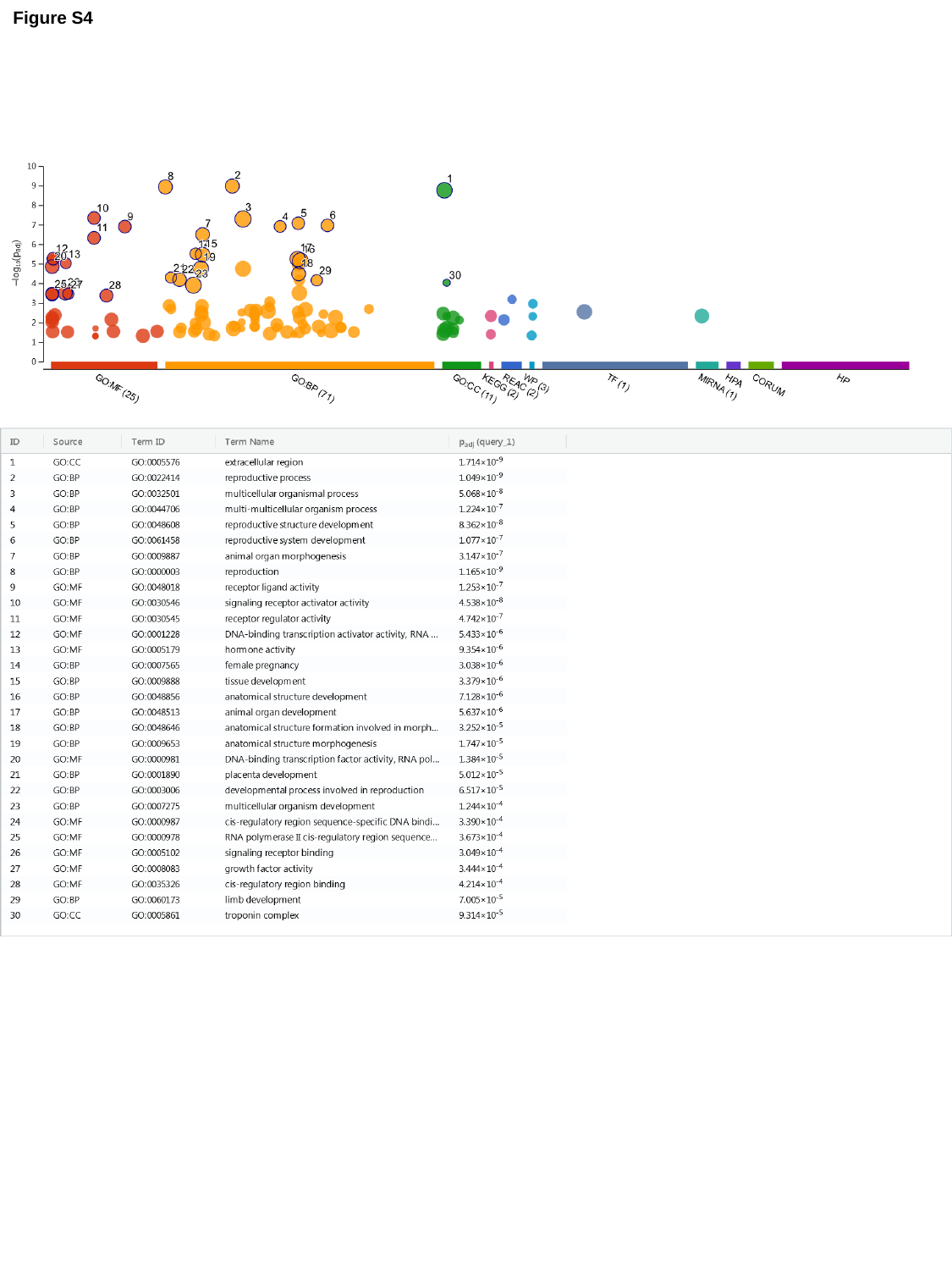

Figure S4

### Slide 5
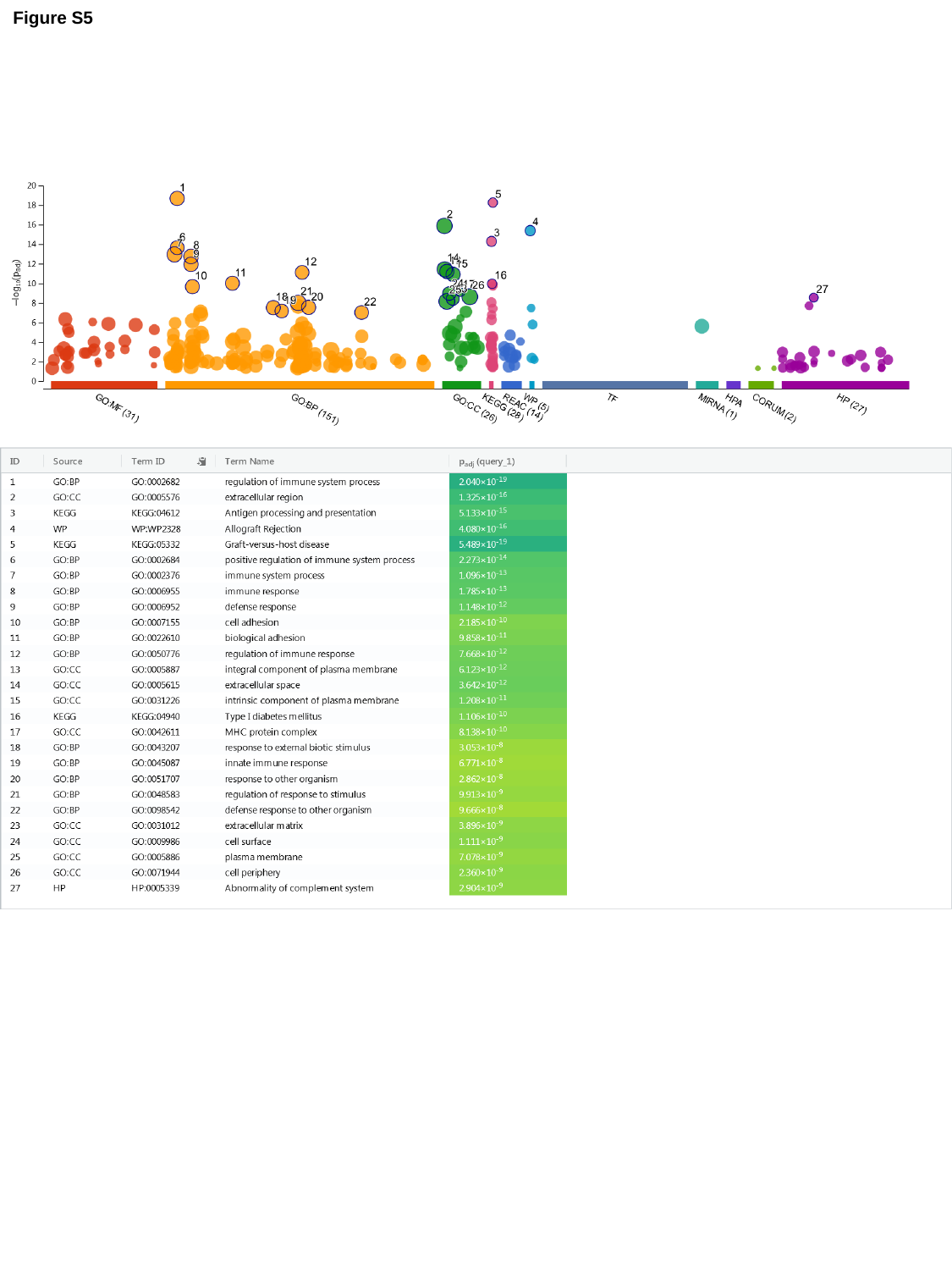

Figure S5

### Slide 6
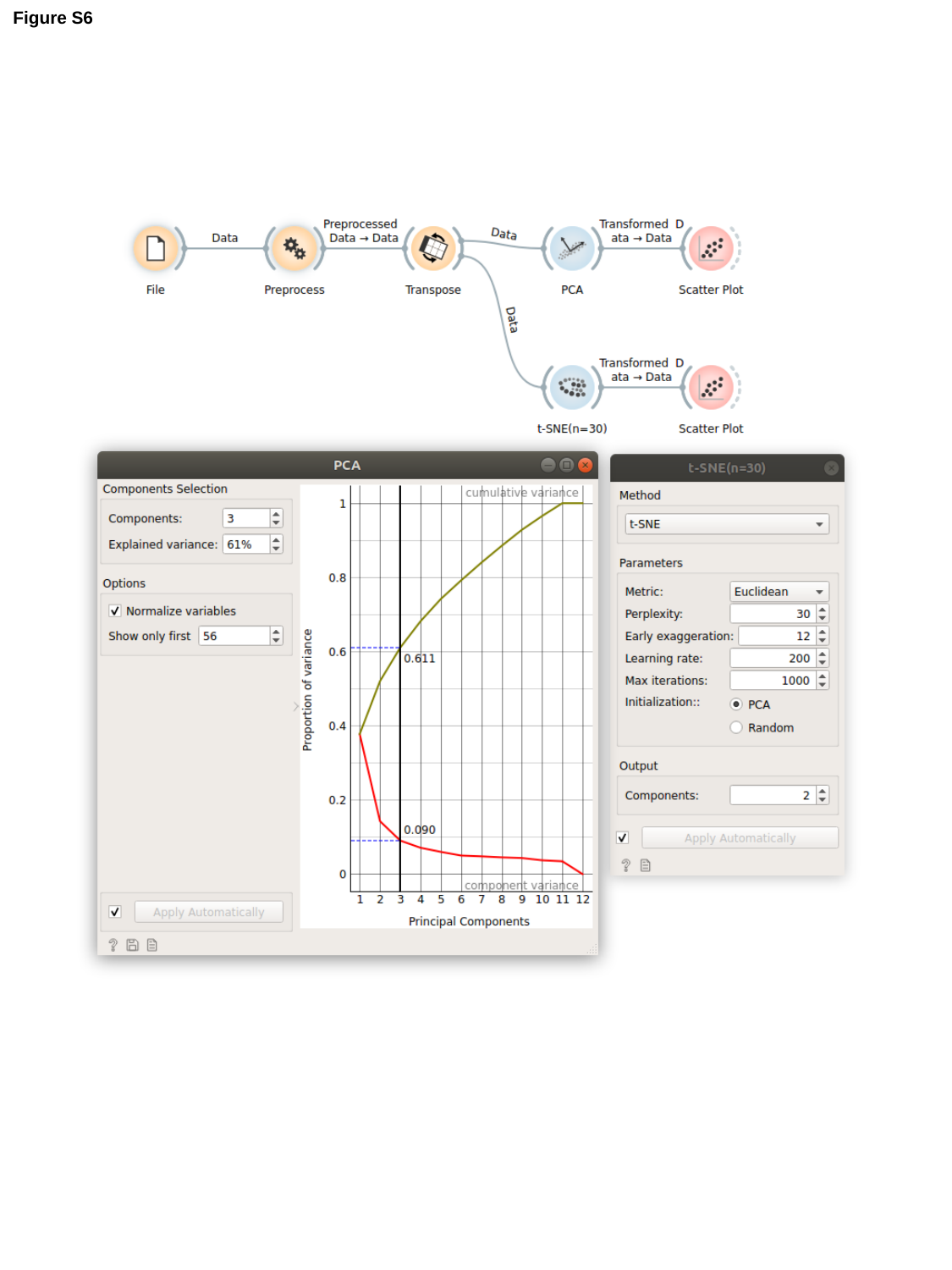

Figure S6

### Slide 7
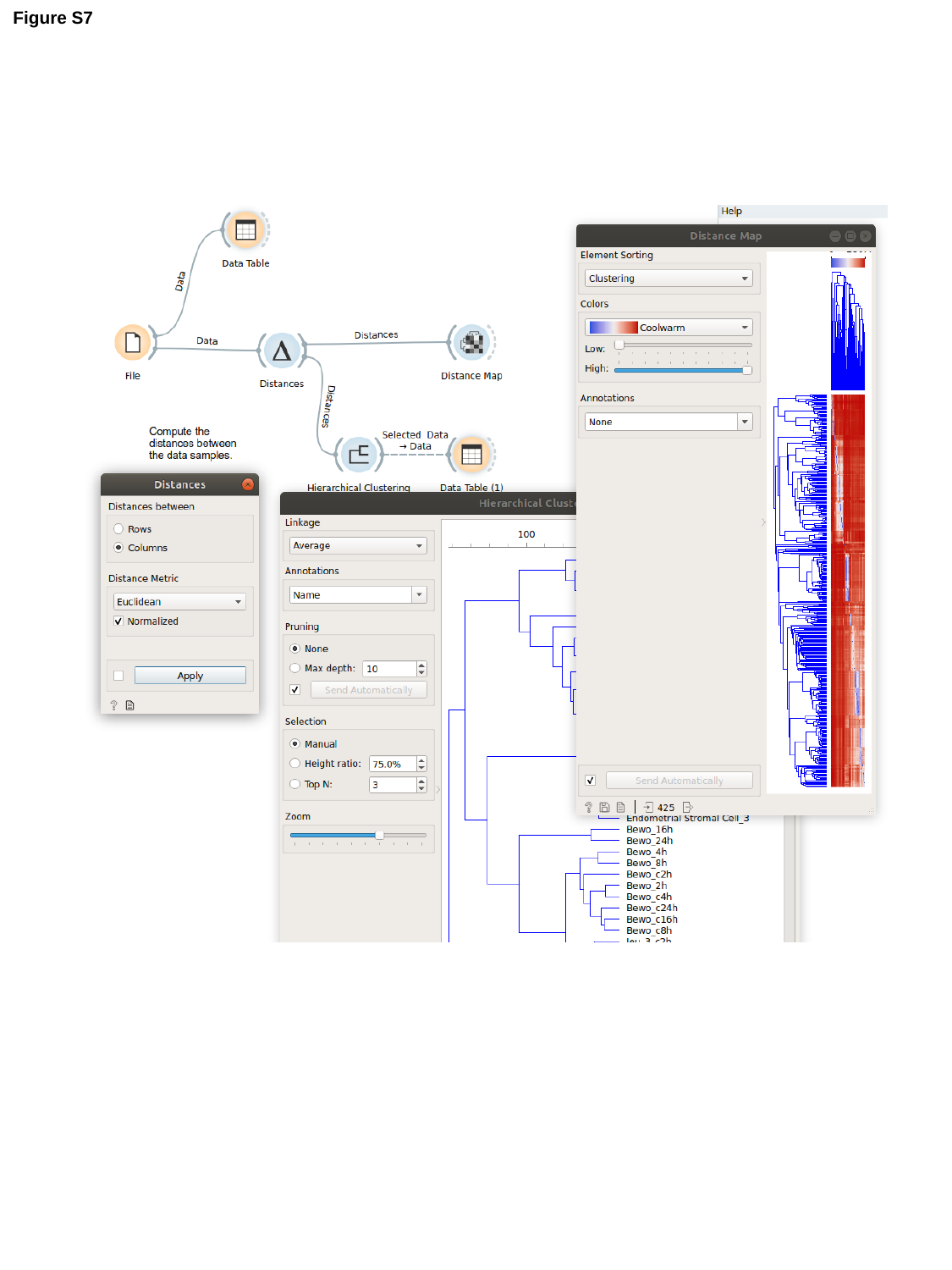

Figure S7

### Slide 8
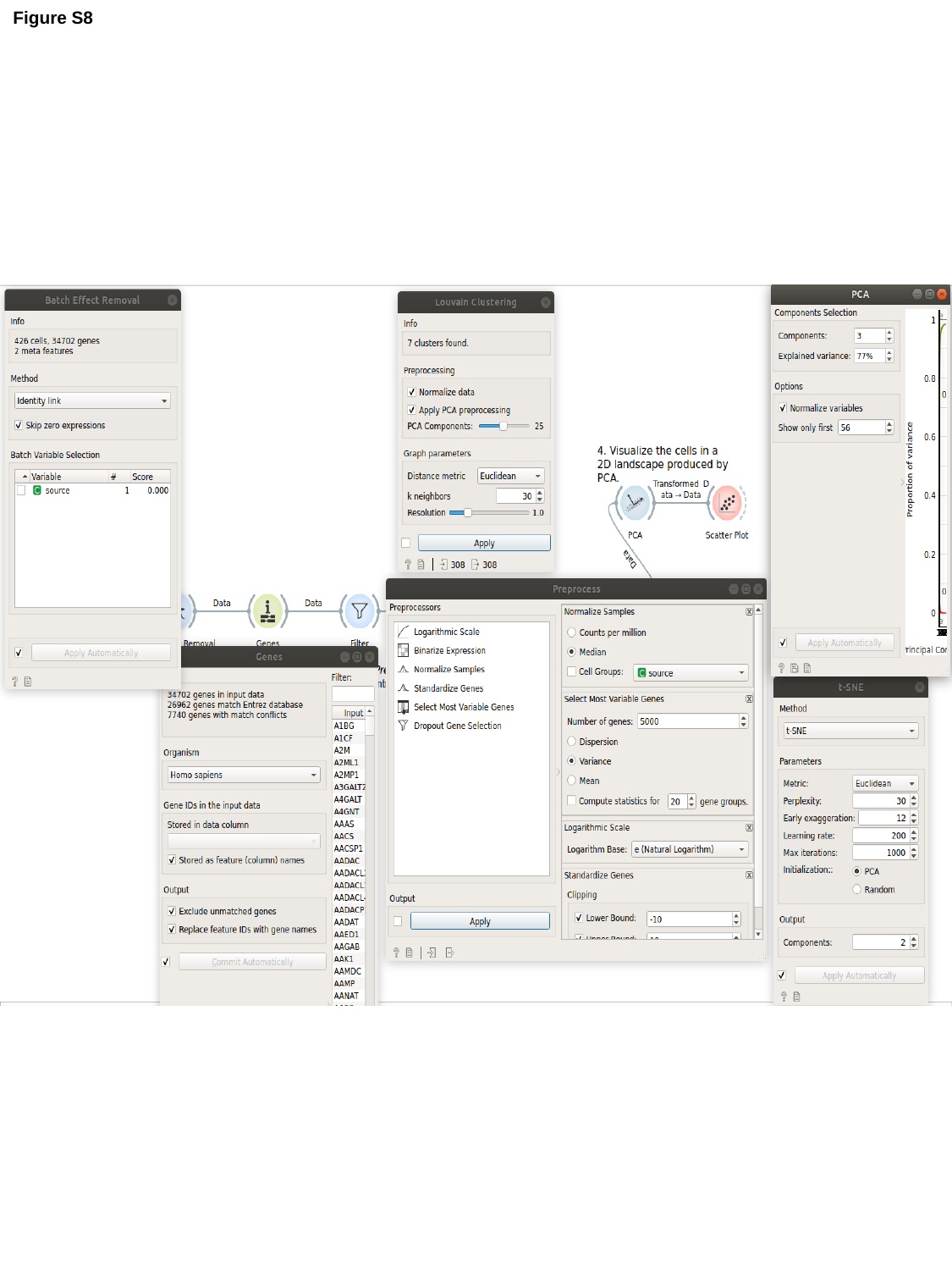

Figure S8

### Slide 9
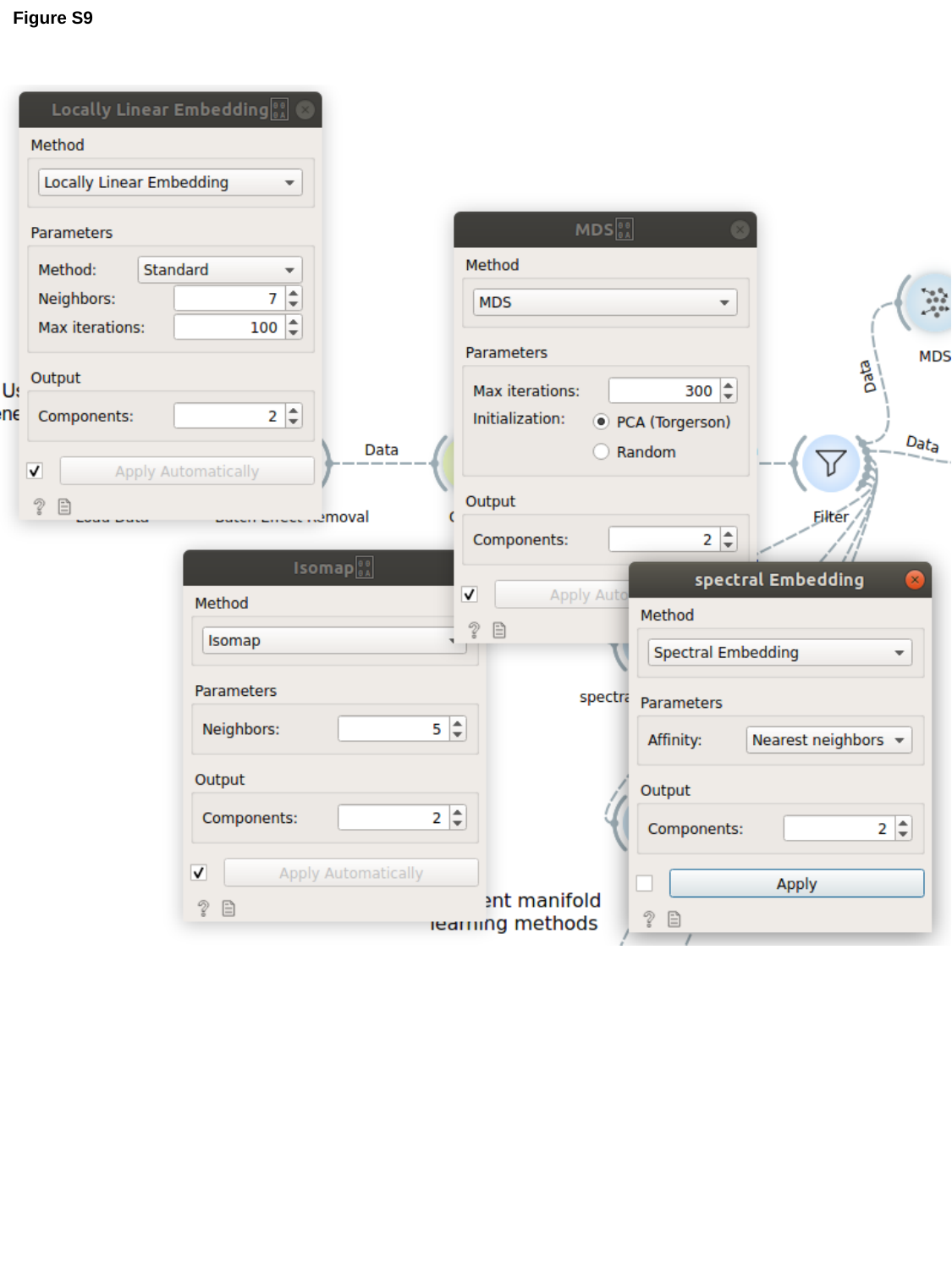

Figure S9

### Slide 10
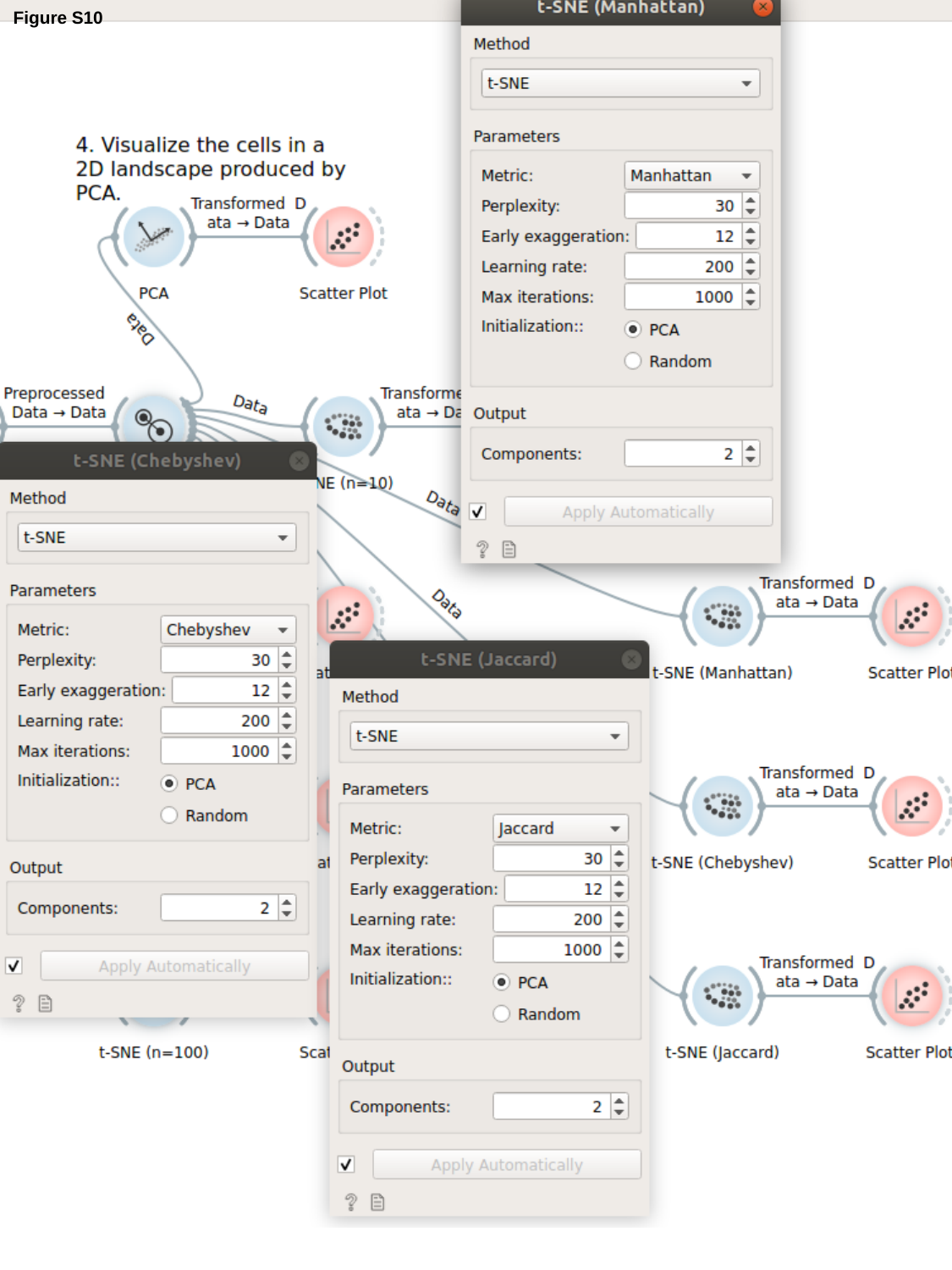

Figure S10
